# Histone Variant H2A.Z Cooperates with EBNA1 to Maintain Epstein-Barr Virus Latent Epigenome

**DOI:** 10.1101/2025.01.28.635203

**Authors:** Josue Leonardo Castro-Muñoz, Davide Maestri, Leena Yoon, Bhanu Chandra Karisetty, Italo Tempera, Paul Lieberman

## Abstract

Chromatin structure plays a central role in the regulation of Epstein-Barr Virus (EBV) latency. The histone variant H2A.Z.1 has been implicated in chromatin structures associated with initiation of transcription and DNA replication. Here, we investigate the functional role of H2AZ.1 in the regulation of EBV chromatin, gene expression and copy number during latent infection. We found that H2A.Z.1 is highly enriched with EBNA1 binding sites at *oriP* and Qp, and to a lesser extent with transcriptionally active CTCF binding sites on the EBV genomes in both Mutu I Burkitt lymphoma (BL) and SNU719 EBV-associated gastric carcinoma (EBVaGC) cell lines. RNA-interference depletion of H2A.Z.1 resulted in the reactivation of viral lytic genes (ZTA and EAD) and increases viral DNA copy numbers in both MutuI and SNU719 cells. H2A.Z depletion also led to a decrease in EBNA1 binding to *oriP* and *Qp*, on the viral episome as well as on oriP plasmids independently of other viral genes and genomes. H2A.Z.1 depletion also reduced peaks of H3K27ac and H4K20me3 at regulatory elements in the EBV genome. In the cellular genome, H2A.Z.1 colocalized with only a subset of EBNA1 binding sites and H2A.Z.1 depletion altered transcription of genes associated with *myc targets* and *mTORC1 signaling*. Taken together, these findings indicate that H2A.Z.1 cooperates with EBNA1 to regulate chromatin structures important for epigenetic programming of the latent episome.

**Importance:** Cellular factors the restrict latent viral reactivation are of fundamental importance. We have found that the cellular histone variant H2A.Z functions in cooperation with the Epstein-Barr Virus (EBV) latency maintenance protein EBNA1 to establish a stable epigenome and restrict lytic cycle reactivation during latency. We show that H2A.Z co-occupies EBNA1 binding sites on the EBV and host genome, and that depletion of H2A.Z leads to robust reactivation of EBV from latency. H2A.Z is important for the function of EBNA1 at the origin of plasmid (oriP) replication and establishing EBV epigenetic marks. H2A.Z binds with EBNA1 at cellular binding sites and controls the expression of cellular genes in the cMyc and mTORC1 pathways that are also implicated in control of EBV latency.

## Introduction

Epstein-Barr virus (EBV) is a highly prevalent human gammaherpesvirus that establishes long-term latent infection in memory B-cells (1–3). EBV is also strongly associated with diverse cancers, including endemic forms of Burkitt’s lymphoma (BL) and nasopharyngeal carcinoma (NPC), numerous other B-lymphomas, NK/T lymphomas and a subtype of gastric carcinomas (GC) (4, 5). In total, EBV is responsible for 1-2% of all human cancers (6).

Most EBV cancers are associated with latent forms of EBV where viral DNA is maintained in tumor cell nuclei as multicopy episomes (7, 8). These episomes are subject to chromatin-associated epigenetic regulation whereby only a few viral genes are expressed, and productive lytic cycle gene expression and replication are largely repressed (9, 10). EBV latency has been categorized into at least four different gene expression programs, called latency types (11). The latency types correlate with host cell and tumor types, although some viral gene patterns can be highly heterogeneous in a tumor population(12, 13). The variable outcome of EBV infection from latent to lytic, or benign to malignant depend on epigenetic changes in viral and host genomes (14). Viral and cellular factors both contribute to the formation and stability of these different epigenetic states.

Central to the epigenetic control of gene expression is the composition of histones and histone variants that assemble as nucleosomes on genomes and gene regulatory elements (15, 16). Histone variants are incorporated into the nucleosomes through histone chaperones and replace canonical histones, conferring a structural and qualitative change to chromatin(17, 18). Histone variants play an important role in various cellular processes such as embryonic development, chromosomal segregation, transcriptional regulation, DNA repair (17, 19). The role of histone variants on the EBV genome is not completely understood. The EBV tegument protein BNRF1 has been found to bind histone chaperone DAXX in complex with H3.3-H4 histones during the early phase assembly of chromatin on the EBV episomes (20, 21). Early studies examining Encode data sets identified the histone variant H2A.Z.1 (subsequently referred to as H2AZ) as potentially enriched at the EBV *oriP* region (22, 23). H2A.Z.1 is more common of two isoforms, the other being H2A.Z.2 (24, 25). H2A.Z family are one of several histone H2A variants that include H2AX and macroH2A (26). H2A.Z.1 is of particular interest since it has been implicated in control of replication origins, DNA repair, centromeric heterochromatin and transcription (activation or repression) (25, 27). H2A.Z.1 is enriched at transcriptional regulatory elements, including promoters, enhancers and insulators, where it is thought to destabilize the core histone and facilitate dynamic chromatin remodeling (27). In addition, overexpression of H2A.Z.1 has been correlated with the development of several cancer types (e.g. pancreas, breast, bladder, melanoma), since its presence in promoter regions favors the expression of genes associated with cell transformation (28–32).

Here we evaluated the role of H2A.Z in the control of EBV latency. We found that H2A.Z is enriched at several sites across the EBV genome with particularly high enrichment around EBNA1 binding sites at *oriP* and *Qp*. We also found that depletion of H2A.Z reduced EBNA1 binding at these viral sites and led to the reactivation of lytic gene expression from latently infected Mutu I Burkitt lymphoma cell lines and SNU719 EBV-associated gastric carcinoma (EBVaGC) cell lines. Our findings suggest that H2A.Z cooperates with EBNA1 binding at both viral and cellular sites to maintain a stable latent infection in EBV infected cancer cells.

## Results

### Histone variant H2A.Z bind in common sites with EBNA1 in EBV genome

We first set out to determine if histone variant H2A.Z is incorporated into the EBV genome. We performed Chromatin Immunoprecipitation followed by sequencing (ChIP-seq) for H2A.Z and EBNA1 in two different EBV latency models, SNU719, an EBVaGC-derived cell line and Mutu I, an EBV^+^ BL-derived cell line. In **Figure 1A** we show the EBV genome from IGV browser with H2A.Z ChIP-seq tracks (magenta) along with EBNA1 ChIP-seq tracks (blue) and Input (grey). For reference, we also provide the CTCF ChIP-seq tracks (green). Analysis of the ChIP-seq peak patterns reveal that H2A.Z has a broad peak covering most of the *oriP* (FR and DS) region and a second large peak at the Qp region. We also observed weaker peaks that colocalized with some CTCF binding sites including those at the LMP1/LMP2 regulatory locus (referred to as CTCF_166_). At the *oriP* locus it is apparent that H2A.Z peak is enriched at both the EBNA1 binding sites at the family of repeats (FR) and dyad symmetry (DS) regions of *oriP* as well as extending across the CTCF binding site at the EBERp located several hundreds of basepairs upstream of the FR and a second CTCF site located just downstream of the DS (**Fig. 1B**). Similarly, the strong H2A.Z peak at Qp is split and centered over the 2 EBNA1 binding sites and the CTCF binding site that is less than 50 bp from the 5’ EBNA1 binding site (**Fig. 1C**). We confirmed by ChIP-qPCR that H2A.Z was enriched at FR, DS, Qp but not at control oriLyt region in SNU719 (**Fig. 1D**) and Mutu I (**Fig. 1E**), as well as in LCL-352 (**Fig. 1F**) and nasopharyngeal carcinoma (NPC) derived C666-1 (**Fig. 1G**). These results indicate that histone variant H2A.Z is enriched on EBNA1 occupied sites in different EBV latency and cell types.

**Figure 1.**
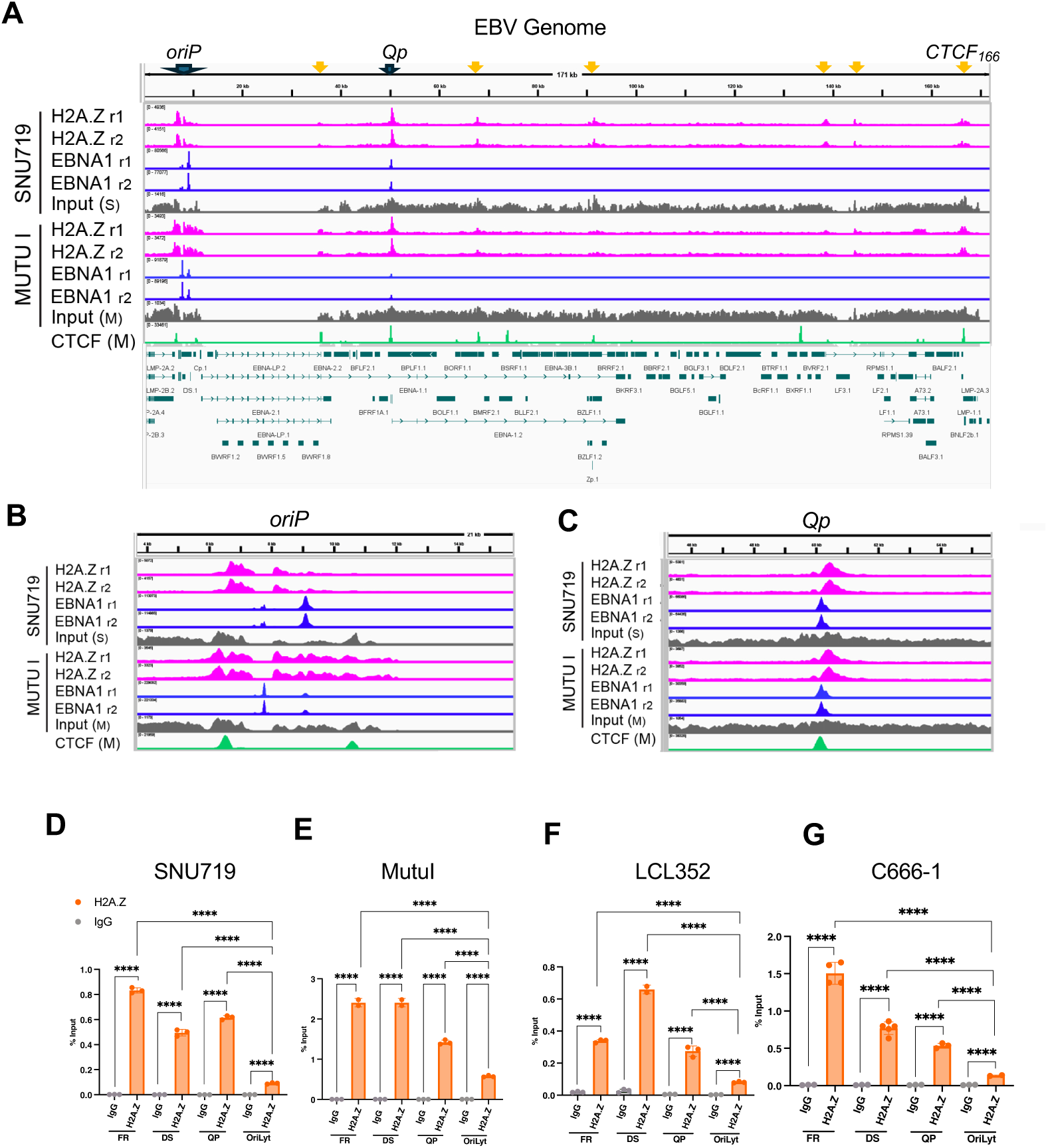
H2A.Z is enriched at EBNA1 and CTCF binding sites in the EBV genome. **(A)** EBV genome visualized using IGV showing ChIPseq tracks for H2A.Z (pink), EBNA1 (blue), and Input controls (grey) for SNU719 (top) or Mutu I (lower). CTCF ChIP-seq for Mutu I is shown in green. EBV genome annotation is provided below. (**B)** magnified ChIP-seq tracks of panel A at EBV *oriP* region. **(C)** Magnified ChIP-seq tracks of panel A at Qp. **(D)** ChIP-qPCR of H2A.Z or IgG control at FR, DS, Qp and *oriLyt* regions of EBV in SNU719, **(E)** Mutu I, **(F)** LCL352 and **(G)** C666-1 cells. **** p<.0001, n=3, student t-test.

#### Knockdown of histone variant H2A.Z disrupts EBV latency to induce lytic cycle gene expression and DNA replication

To determine if H2A.Z enrichment on the EBV latent episome has any functional role, we depleted H2A.Z by RNA interference knock-down. We assayed two independent small hairpin RNA (shRNA) using lentivirus transduction in Mutu I cells and validated that these efficiently knocked down H2A.Z by Western blot (**Fig. 2A**). Western blot analysis revealed that H2A.Z knock-down partially reduced EBNA1 expression and more substantially increased lytic cycle BZLF1 transcriptional activator ZTA relative to actin control (**Fig. 2A**). RT-qPCR showed that both shRNAs for H2A.Z induced significant levels of EBV lytic transcripts for ZTA, as well as EA-D (**Fig. 2B**). DNA qPCR comparing viral *oriLyt* to cellular GAPDH ratios showed a significant increase in viral DNA copy number after transduction with both shH2A.Z relative to shControl lentivirus transduced Mutu I cells (**Fig. 2C**). We next tested the effects of siRNA depletion of H2A.Z in SNU719. We used an siRNA pool targeting H2A.Z (ON-TARGETplus Smartpool) or non-targeting control siRNA in these EBVaGC cells since these are more transfectable than Mutu I. Western blot demonstrated that H2A.Z was efficiently depleted by siH2A.Z (**Fig. 2D**). Similar to our findings in Mutu I, H2A.Z depletion led to a modest loss of EBNA1 protein, and a large increase in the lytic protein EA-D (**Fig. 2D**). RT-qPCR showed that siH2A.Z induced both Zta and EA-D transcripts (**Fig. 2E**) and an increase in viral DNA copy number as measured by qPCR (**Fig. 2F**). While the DNA amplification in SNU719 is not as robust as in Mutu I cells, the general trend towards lytic gene activation and viral DNA amplification were similar in both cell types. These findings indicate that H2A.Z depletion leads to a disruption of EBV latency with an increase in lytic cycle gene transcripts and viral DNA copy number.

**Figure 2.**
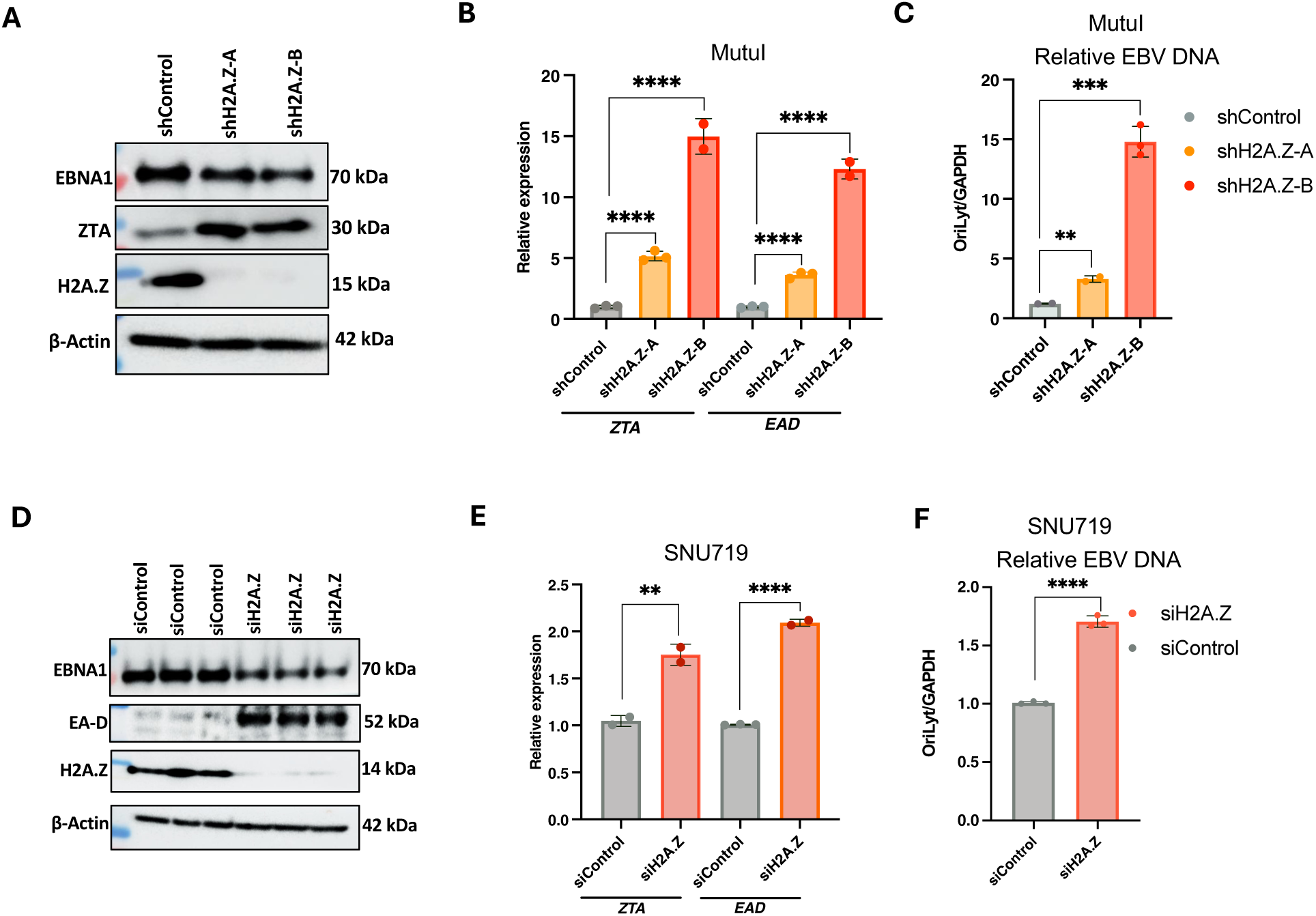
H2A.Z knock-down activates EBV lytic gene expression. **(A)** Mutu I cells transduced with lentivirus expressing shControl, shH2A.Z-A or shH2A.Z-B were selected for puromycin resistance for 7 days and then assayed by Western blot for EBNA1, ZTA, H2A.Z or β-Actin. **(B**) H2A.Z knock-down in Mutu I cells were assayed by RT-qPCR for *ZTA* and *EA-D* mRNA expression. **(C)** EBV DNA copy number measured by qPCR comparing EBV *oriLyt* DNA relative to cellular GAPDH DNA for samples as described in panel A. **(D)** SNU-719 cells were transfected with siControl or siH2A.Z and assayed by Western blot for EBNA1, EA-D, H2A.Z, and β-Actin. **(E)**. SNU719 cells treated as described for panel D were assayed by RT-qPCR for *ZTA* and *EA-D* mRNA expression. **(F)** EBV DNA copy number measured by qPCR comparing EBV *oriLyt* DNA relative to cellular GAPDH DNA for samples as described in panel D. ** p<.01, *** p<.001, **** p<.0001 student t-test, n=3.

#### Depletion of H2A.Z reduces EBNA1 binding to the EBV genome

We next examined the effects of H2A.Z knockdown on EBNA1 binding to the EBV genome (**Fig. 3**). We used ChIP-qPCR to assay EBNA1 binding to the FR and Qp, and as a negative control *oriLyt* (**Fig. 3A and B**). We observed a substantial reduction (∼2-3 fold) in EBNA1 binding at both FR and Qp in Mutu I (**Fig. 3A**) and SNU719 (**Fig. 3B**). As expected, EBNA1 did not bind *oriLyt* demonstrating the specificity of EBNA1 binding to EBV genome. These findings suggest that H2A.Z is important for EBNA1 binding to the viral episome during latent infection.

**Figure 3.**
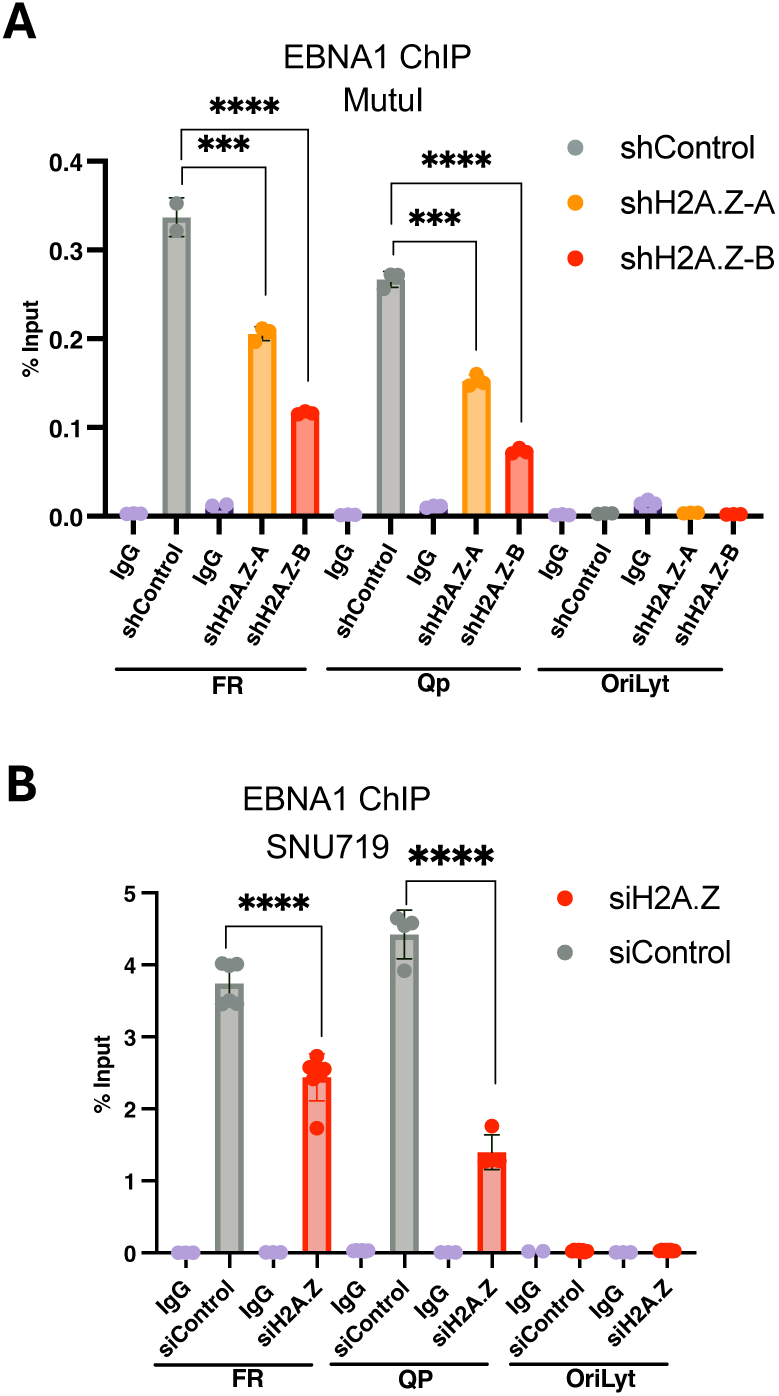
H2A.Z knock-down reduces EBNA1 binding to EBV genome. **(A)** Mutu I cells transduced with shControl, shH2A.Z-A, or shH2A.Z-B were analyzed by Western blot for EBNA1, H2A.Z, or β-Actin. **(B)** ChIP-qPCR for EBNA1 or IgG control at FR, Qp and *oriLyt* regions of EBV in Mutu I cells transduced with shControl, shH2A.Z-A, or shH2A.Z-B. **(C)** SNU-719 cells transfected with siControl or siH2A.Z were analyzed by Western blot for EBNA1, H2A.Z, or β-Actin. **(D)** ChIP-qPCR for EBNA1 or IgG control at FR, Qp and *oriLyt* regions of EBV in SNU719 cells treated with siControl or siH2A.Z. *** p<.001, **** p<.0001 student t-test, n=3.

#### H2A.Z knock-down reduces EBNA1-binding and oriP-dependent plasmid DNA replication

Since H2A.Z knock-down can induce viral lytic cycle gene expression and DNA replication in EBV positive cell lines, it is possible that this indirectly inhibits EBNA1 DNA binding to EBV episomes. To eliminate this concern, we assayed the effect of H2A.Z knockdown in EBV-negative 293T cells transfected with a self-replicating plasmid expressing FLAG-EBNA1 and containing the *oriP* replicon (**Fig. 4**). We validated that H2A.Z could be efficiently depleted in 293T cells transfected with H2A.Z siRNA pool (**Fig. 4A**). ChIP-qPCR was used to assay H2A.Z and EBNA1 binding after siControl and siH2A.Z. H2A.Z bound significantly to the FR and DS regions of the *oriP* plasmid, and to a lesser extent the Ampicillin (Amp) gene, located ∼4 kb away from *oriP* on the same plasmid DNA. As expected, siRNA depletion of H2A.Z led to a significant reduction of its binding to *oriP* (**Fig. 4B**). EBNA1 bound efficiently to the DS and FR, but not to the Amp gene of the *oriP* plasmid in siRNA control cells (**Fig. 4C**). However, in H2A.Z depleted cells, EBNA1 binding was reduced ∼7 fold at the DS and FR elements (**Fig. 4C**). We also assayed the relative DNA copy number of the *oriP* plasmid using qPCR for either the DS or FR element relative to cellular GAPDH DNA (**Fig. 4D**). Using both markers of the *oriP* plasmid, we observed a significant decrease in the *oriP* plasmid copy number after siRNA depletion of H2A.Z relative to siControl (**Fig. 4D**). Taken together, these findings indicate that H2A.Z facilitates the binding of EBNA1 to *oriP* and consequently, the EBNA1-dependent DNA replication/episome maintenance of the *oriP*-containing plasmid.

**Figure 4.**
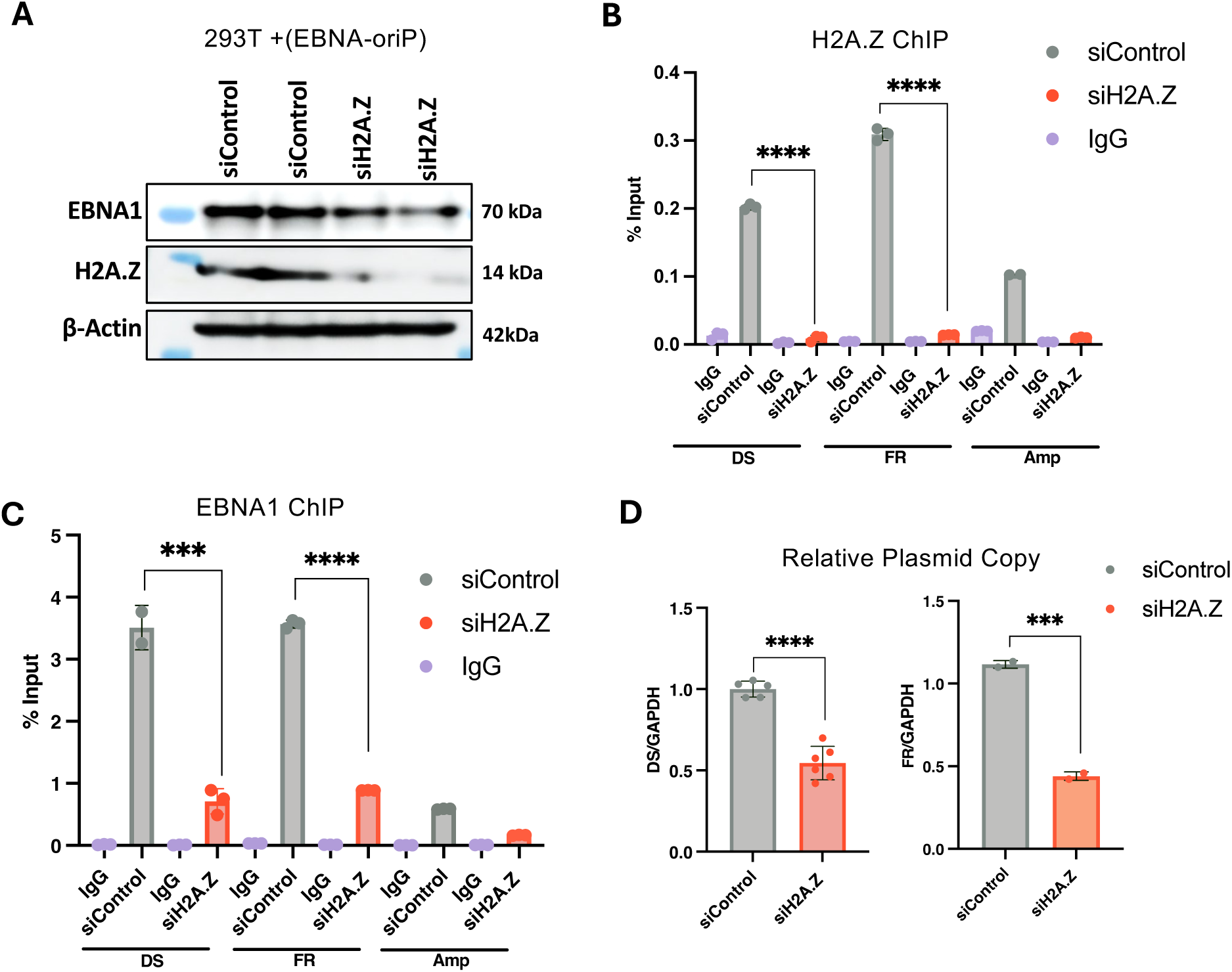
H2A.Z depletion reduces EBNA1 binding to OriP in EBV^-^ 293T cells. 293T cells transfected with oriP plasmid expressing EBNA1 were co-transfected with siControl or siH2A.Z. **(A)** Assayed by Western blot for expression of EBNA1, H2A.Z or β-Actin. **(B)** ChIP-qPCR for H2A.Z or control IgG at the DS, FR or AMP regions of the *oriP* plasmid in transfected 293T cells. **(C)** ChIP-qPCR for EBNA1 or control IgG at the DS, FR, or AMP region of the oriP plasmid in transfected 293T cells and **(D)** *OriP* plasmid copy number determined by qPCR using primers for *oriP* elements DS (left) or FR (right) comparing siControl or siH2A.Z in 293T cells. *** p<.001, **** p<.0001 student t-test, n=3.

#### EBNA1 enhances H2A.Z binding at *oriP*

To determine if EBNA1 contributes to the enrichment of H2A.Z occupancy at *oriP*, we performed ChIP-qPCR assays with an *oriP*-containing plasmid lacking EBNA1 (pHEBO) in 293T cells co-transfected with either empty vector or with pCMV-FLAG-EBNA1 (**Fig. 5**). Expression of FLAG-EBNA1 was validated by Western blot (**Fig. 5A**) and its efficient binding to DS and FR was shown by ChIP-qPCR (**Fig. 5B**). EBNA1 bound weakly to the Amp regions, as may be expected since this is only ∼4 kb from the *oriP* element in the same plasmid. ChIP-qPCR for H2A.Z revealed a significant increase (∼3-5 fold) of binding at the DS and FR in the presence of EBNA1 (**Fig. 5C**). In contrast, H2A.Z binding slightly decreased at the Amp gene. These findings indicate that EBNA1 enhances the binding of H2A.Z at EBNA1 binding sites in *oriP*.

**Figure 5.**
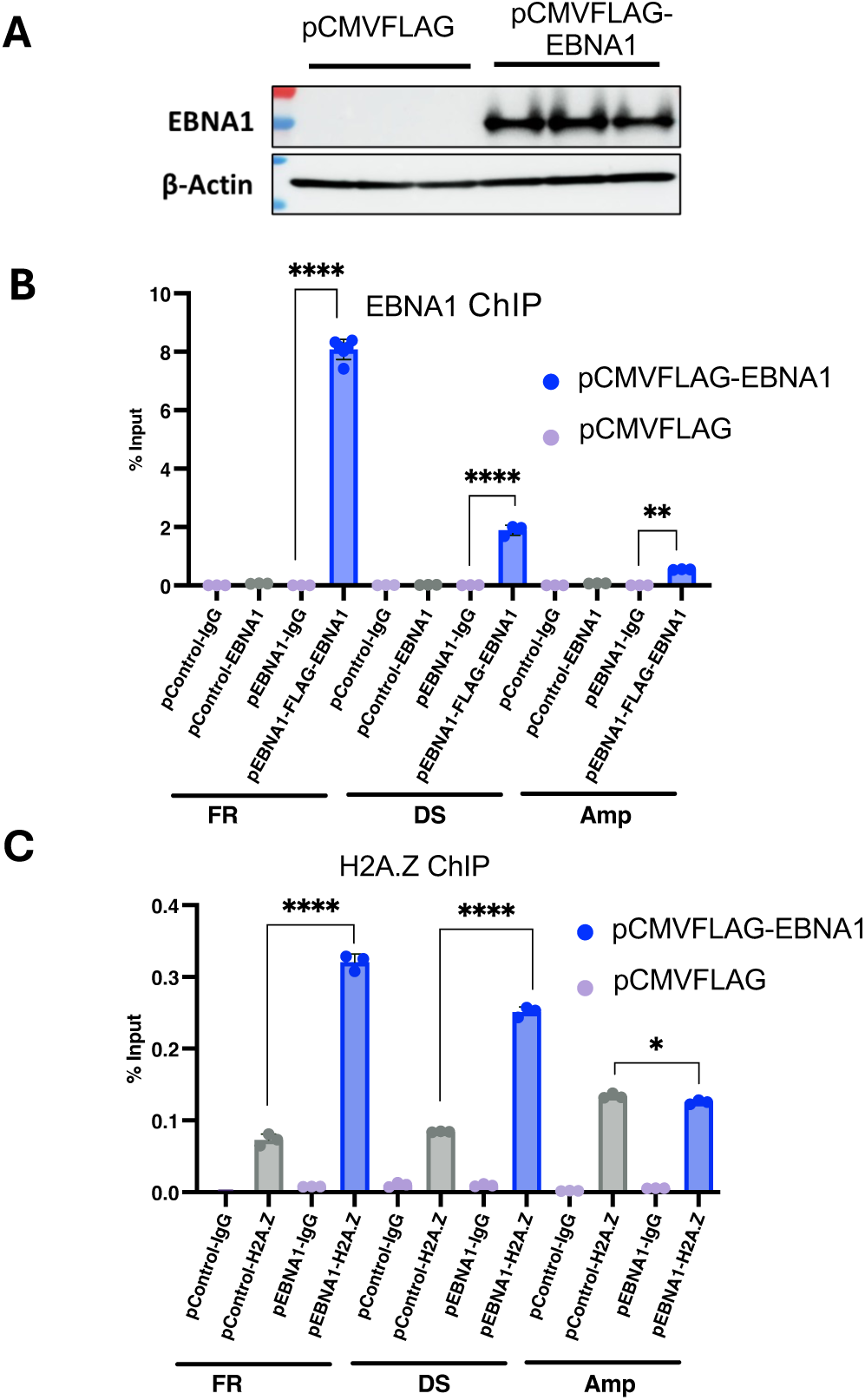
EBNA1 increases H2A.Z enrichment at *oriP*-containing plasmids in 293T cells. 293T cells were transfected with OriP replicon (pHEBO), plasmid expressing FLAG-EBNA1 (pCMVFLAG-EBNA1) and plasmid Control (pCMVFLAG). **(A)** Western blot shows FLAG-EBNA1 and β-Actin protein levels in 3 biological replicates. **(B)** ChIP-qPCR for FLAG-EBNA1 or control IgG at the DS, FR or AMP regions in transfected 293T cells. **(C)** ChIP-qPCR for H2A.Z or control IgG at the DS, FR, or AMP region in transfected 293T cells with FLAG-EBNA1. ***p<.001, ****p<.0001 student t-test, n=3.

#### H2A.Z differentially affects CTCF binding and histone modifications at regulatory sites in the EBV genome

ChIP-seq data showed that H2A.Z colocalized with several CTCF binding sites on the EBV genome, including those at the Qp and LMP1/LMP2 locus (CTCF_166_) (**Fig. 1A-C**). We therefore assayed the effect of H2A.Z depletion on CTCF binding to Qp and LMP1/LMP2 locus CTCF sites. ChIP-qPCR for CTCF revealed that H2A.Z knock-down modestly increased (∼1.3 fold) CTCF binding at Qp and CTCF_166_ binding sites. No CTCF binding was detected at the oriLyt region which was used as a specificity control. The observed increase binding of CTCF contrasts with the decreased binding by EBNA1, suggesting that H2A.Z has opposing effects on these two different DNA binding factors.

H2A.Z is also known to contribute to both active and repressive cellular chromatin structures. To explore the potential role of H2A.Z on histone modifications associated with either active or repressive chromatin on the EBV genome. We evaluated the enrichment of four histone marks at two regions of the EBV genome that have different epigenetic regulation, namely Qp, which is transcriptionally active in type I latency and *oriLyt,* which is silenced for lytic DNA replication in SNU719 cells. Two histone marks associated with transcriptional activation such as H3K27ac and H3K40me3, and two histone marks associated with repression like H3K27me3 and H4K20me3 were evaluated (**Fig. 6B and C**). We first show that H3K27ac was more enriched in Qp region (**Fig. 6B**) while H4K20me3 was more enriched in *oriLyt* region (**Fig. 6C**). We next analyzed the effects of H2A.Z knock-down on these histone marks. We found that H2A.Z knock-down led to a loss of H3K27ac at Qp and H4K20me3 at *oriLyt* (**Fig. 6D and E**). These results indicate that H2A.Z contributes to the formation of histone modifications at key regulatory elements throughout the EBV epigenome.

**Figure 6.**
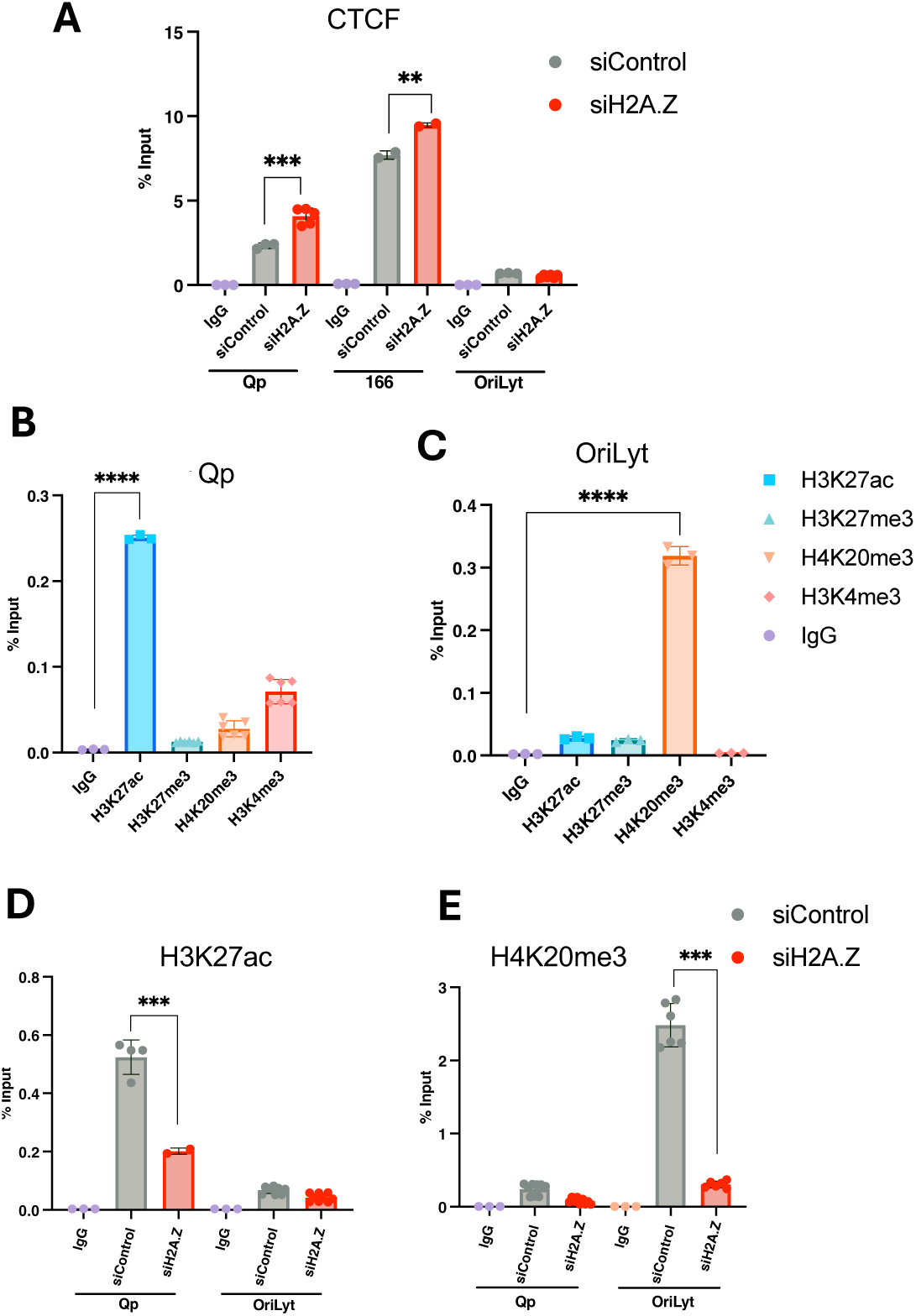
H2A.Z depletion alters histone modifications on the EBV genome. **(A)** ChIP-qPCR for CTCF or control IgG for DS, CTCF_166_, or *oriLyt* regions of EBV genome in SNU-719 cells treated with siControl or siH2A.Z. **(B-C)** ChIP-qPCR for H3K27ac, H3K27me3, H4K20me3, H3K4me3 or IgG control at the Qp region (panel C) or the *oriLyt* region (panel D) in untreated SNU719 cells. **(D-E)** ChIP-qPCR for H3K27ac (panel E) or H4K27me3 (panel F) or control IgG at Qp or *oriLyt* regions of EBV in SNU719 cells treated with siControl or si-H2A.Z. *** p<.001, **** p<.0001 student t-test, n=3.

#### Histone variant H2A.Z and EBNA1 are able bind in common sites in cellular genome

We next examined the binding patterns of EBNA1 and H2A.Z in the cellular genomes of these EBV^+^ cells. We analyzed genome-wide ChIP-seq binding of H2A.Z and EBNA1 in Mutu I and SNU719. In **Figure 7A** we show heatmap alignments of all EBNA1 binding sites compared to H2A.Z binding sites which revealed very distinct patterns in both cell lines, with EBNA1 peaks being very narrow while H2A.Z peaks appear to be broader, ranging from 100 to 1000 bp. More specifically, in Mutu I cells we identified 6819 regions enriched with H2A.Z, of which 5.7% were in promoter regions and 94.3% was in intergenic regions, while EBNA1 was enriched in 1102 regions, of which 2.1% was in promoter regions and 97.9% was in distal intergenic regions (**Fig. 7B**, upper panel). In SNU719, H2A.Z was enriched in 8796 regions and only 9.5% was in promoter regions and 90.6% was in distal intergenic regions, while EBNA1 was enriched in 968 regions, of which 3.4% was in promoter regions and 96.6% was in intergenic regions (**Fig. 7B**, right). Venn diagrams show that H2A.Z and EBNA1 binding overlaps in approximately 700 regions in both cell lines (**Fig. 7B**, lower panel). Overlap of EBNA1 and H2A.Z binding sites were also shown by heatmap analysis showing a small fraction of H2A.Z and EBNA1 sites directly overlap across the cellular genome (**Fig. 7C**). Examination of several previously documented EBNA1 peaks, including ADA and IL6R show interesting patterns of H2A.Z overlap and interaction (**Fig. 7D**). At the ADA locus, where EBNA1 binds to an upstream enhancer (33), the binding pattern resembles that at *oriP* where the H2A.Z peak is broadly spread over EBNA1 peak and bounded by CTCF sites on each end. At the IL6R gene (34), EBNA1 is bound to the TSS while H2A.Z is bounded by EBNA1 at the 5’ end but then distributes more broadly throughout the body of the transcribed gene (**Fig. 7D**, middle paneI). At the SELENOK locus, EBNA1 and H2A.Z overlap at the TSS and H2A.Z is enriched at a downstream element linked to the EBNA1 site as detected by Genehancer (35) (**Fig. 7D**, lower panel). These observations suggest that EBNA1 and H2A.Z functionally interact at a subset of cellular genes bound by EBNA1 and important for EBV infection and cellular transformation.

**Figure 7.**
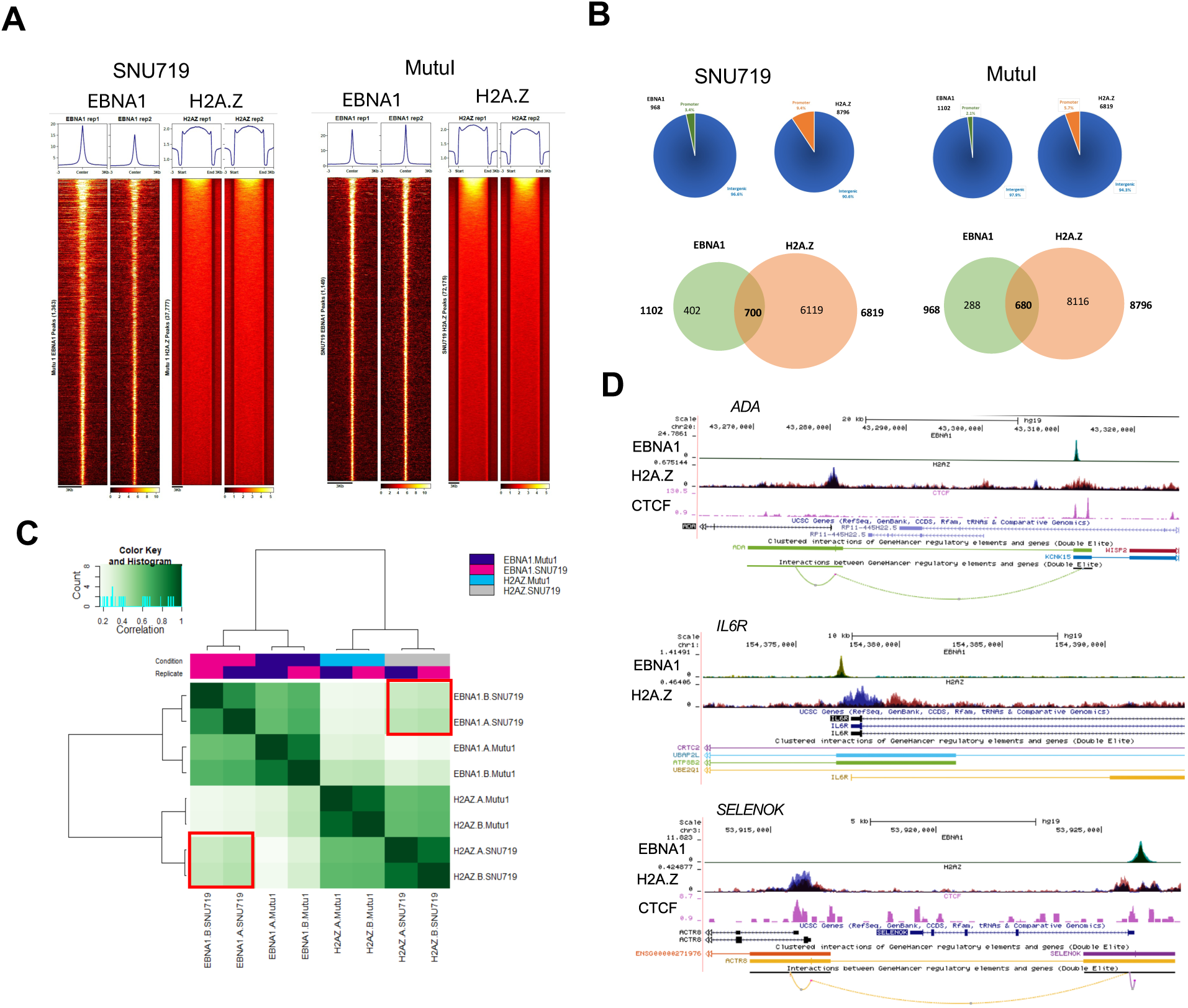
Host genome binding patterns of EBNA1 and H2A.Z. **(A)** Heat map showing the different patterns of ChIP-seq peak distribution for EBNA1 (left) or H2A.Z (right) in SNU719 (left group) or Mutu I (right group). **(B)** Pie charts and Venn diagrams showing the distribution of EBNA1 and H2A.Z ChIP-seq peaks across the cellular genome. Pie chart shows distribution between promoter (green or orange) and intergenic (blue) regions. Venn diagrams show the overlap of peaks between EBNA1 and H2A.Z in SNU719 or Mutu I cells. **(C)** Heatmap showing the overlap correlations of ChIP-seq peaks for H2A.Z and EBNA1 in SNU719 and Mutu I cells. **(D)** ChIP-seq tracks for EBNA1, H2A.Z and CTCF shown at the ADA (top) or IL6R (middle) or SELENOK (bottom) gene loci. Combined tracks for H2A.Z show Mutu I (blue) and SNU719 (red). Gene transcripts and GeneHancer interactions are shown in below each set of tracks.

#### Loss H2A.Z deregulates cellular gene expression

We next asked whether H2A.Z depletion alters host cell genes that are important for EBV infection and cellular transformation. To analyze cellular gene expression after the H2A.Z knockdown in gastric cancer we performed RNA-seq. Data analysis was performed considering a p-value of ≤0.05 as significant, we got that 1414 genes were deregulated after the knockdown. Heatmap analysis identifies clusters of 693 genes that were down-regulated and 721 genes up-regulated (**Fig. 8A**). Gene Ontology analysis of these differentially regulated genes identified the pathways of MYC targets and mTORC1 signaling as the most significantly affected pathways (**Fig. 8B**). Gene set enrichment analysis (GSEA) showed that genes associated with MYC and mTORC1 were downregulated after H2A.Z knockdown (**Fig. 8C**). Interestingly, loss of either MYC or mTORC1 have been found to promote lytic reactivation of EBV (36, 37). These results indicate that H2A.Z can regulate host genes associated with MYC and mTORC1 pathways that are known to regulate EBV gene expression.

**Figure 8.**
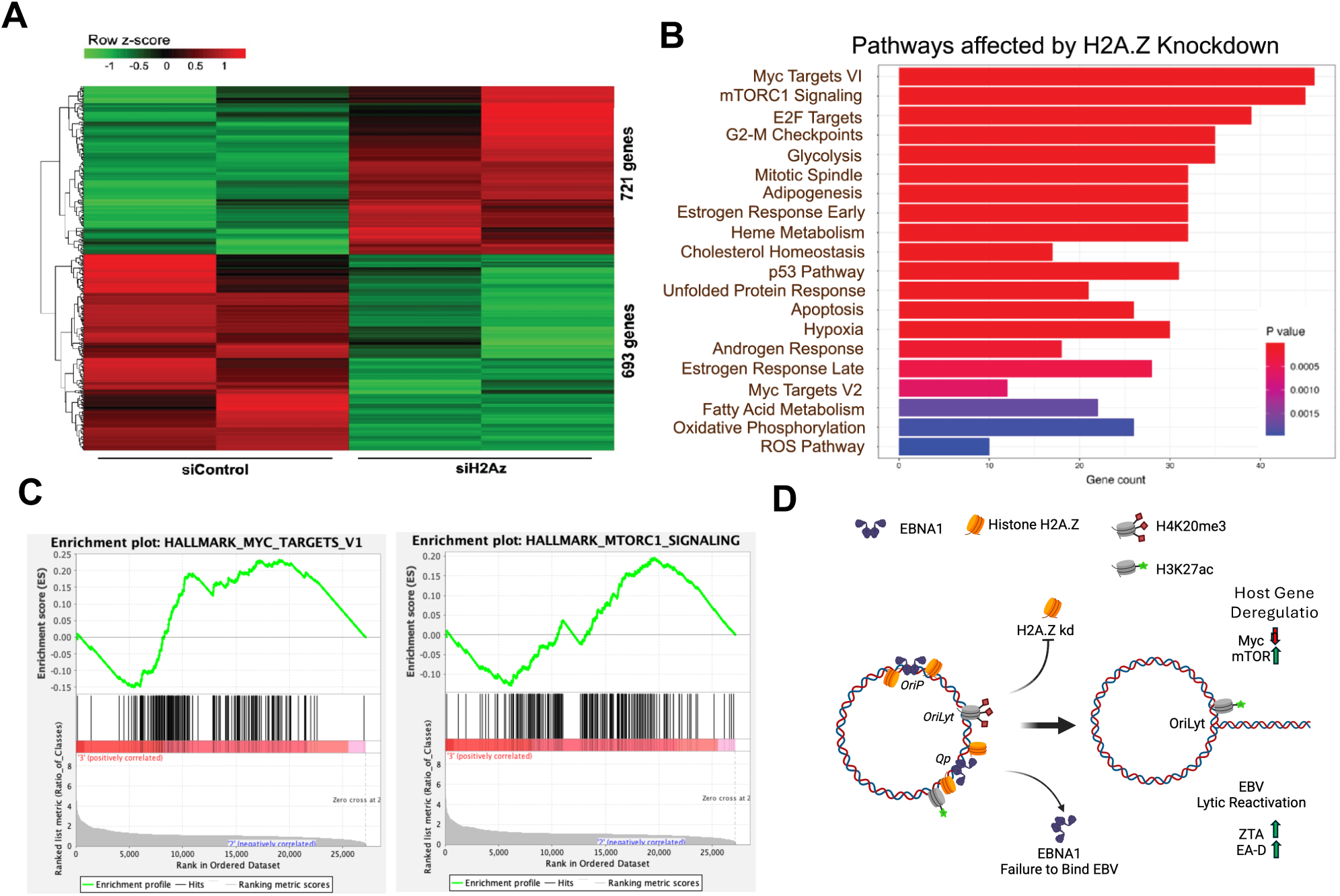
RNAseq analysis of H2A.Z knock-down in SNU719 cells. **(A)** Hierarchical clustering of differentially regulated genes in SNU-719 cells treated with siControl or siH2A.Z. **(B)** Reactome Pathway analysis of genes differentially regulated in SNU-719 cells treated with siH2A.Z or siControl. **(C)** Gene enrichment analysis for MYC Targets (left panel) or PI3K_AKT_MTOR_Signaling (right panel) for RNAseq data set described in panel A. **(D)** Schematic summary of H2A.Z regulation of EBV latency through cooperative binding with EBNA1 and control of c-myc and mTOR pathways.

## Discussion

The gene expression programs that determine EBV latency are governed in large part by chromatin structures and their epigenetic programming (7, 38, 39). Here, we show that the histone variant H2A.Z is enriched at EBNA1 and several CTCF binding sites in the viral genome, and that depletion of H2A.Z leads to the disruption of EBV latency and the reactivation of viral lytic gene expression. We also found that EBNA1 binding and multiple histone modifications are reduced upon H2A.Z knock-down, while CTCF binding increases slightly. Further, H2A.Z enrichment at *oriP* required EBNA1. We found that H2A.Z, which can have both broad and sharp peaks across the cellular genome, overlaps with many EBNA1 binding sites. Finally, we show that depletion of H2A.Z perturbs expression of genes associated with MYC and mTORC regulated pathways, which have been implicated in the control of EBV latency. These findings indicate that H2A.Z plays a viral-specific and global role in maintaining the latent state of the EBV epigenome (**Fig. 8D**).

H2A.Z is a highly conserved histone variant that is essential for organismal development and has been implicated in numerous functional processes, including initiation of DNA replication and RNA transcription, chromosome segregation, response to DNA damage and chromosome structural organization including DNA-DNA looping (24, 27, 40, 41). Nucleosomes containing H2A.Z are thought to be less stable, and therefore contribute to the accessibility and plasticity of chromatin structures (42). We found that H2A.Z was most enriched at regions overlapping EBNA1 binding sites at *oriP* and Qp on the EBV genome. These EBNA1 sites represent an origin of DNA replication (*oriP*) and a transcriptional initiation site for EBNA1 mRNA (Qp). Each of these sites are adjacent to CTCF binding sites and are known to form DNA-DNA loops between each other and other CTCF binding sites in the EBV genome (43, 44). The localization of H2A.Z at oriP was found to be dependent on EBNA1 binding (Fig. 4).

How might EBNA1 facilitate H2A.Z incorporation? Proteomic analysis of EBNA1 interaction partners failed to identify H2A.Z by co-immunoprecipitation (45). Thus, it is unlikely that EBNA1 has a strong, direct interaction with H2A.Z. H2A.Z has been found to localize to cellular origins of DNA replication and facilitate origin licensing in mammalian cells (46, 47). The origin recognition complex 1 (ORC1) protein has been found to have intrinsic histone remodeling activity that can favor H2A.Z loading (47). EBNA1 is known to interact with components of ORC and facilitate its recruitment to the DS element of oriP (48–50). Thus, EBNA1-ORC interactions may facilitate H2A.Z loading at oriP (47). EBNA1 is known to alter nucleosome structure (51), and can also interact with other host proteins that may influence H2A.Z loading, such as nucleosome assembly protein 1 (NAP1) (52, 53). We have previously proposed that EBNA1 has chromatin pioneering activity (33), and our findings here indicate that H2A.Z greatly facilitates EBNA1 binding to chromatinized DNA in vivo, suggesting these proteins may work coordinately to invade and re-organize chromatin at EBNA1 targeted sites.

H2A.Z localization is thought to be regulated by specific chaperones and eviction factors (24, 54). Two complexes that deposit H2A.Z into chromatin are the p400/Tip60/NuA4 complex (55, 56) and the snf2-related CREBBP activator protein (SRCAP) complex (57, 58). H2A.Z can also be actively evicted by chromatin remodeling factors, including Ino80 (59, 60) and the acidic nuclear phosphoprotein 32 family member E (ANP32E) of the p400/Tip60/NuA4 complex (61, 62). Future studies will be required to determine if these factors contribute to the localization of H2A.Z at EBNA1 sites on the EBV and cellular genomes. H2A.Z has also been reported to function in chromatin loop formation through favoring the enhancer RNA (eRNA) expression and facilitating interactions between enhancers and promoters (40). CTCF (CCCTC-binding factor) is also well-established in forming DNA loops, and previous studies have shown that CTCF mediated loops in the EBV genome can also involve contacts with CTCF and EBNA1 bound regulatory elements in *oriP* and Qp (43). We found that H2A.Z knockdown had no effect on CTCF protein levels, but trended to increasing CTCF binding at several colocalized sites across the EBV genome . These findings suggest that H2A.Z may antagonize or compete with CTCF binding at these highly dynamic sites. Future studies will be required to determine if the H2A.Z directly regulates CTCF binding or chromosome 3D structure including DNA loops between CTCF binding sites (25).

Depletion of H2A.Z from EBV infected cells resulted in the disruption of latency and the expression of EBV lytic cycle genes. Since H2A.Z depletion leads to the loss of EBNA1, it is possible that loss of EBNA1 binding destabilizes the latent chromosome to unleash lytic cycle reactivation. Alternatively, it is possible that H2A.Z is required for repression of lytic cycle genes. H2A.Z binding at CTCF sites upstream of BZLF1 could also play in regulating lytic reactivation. H2A.Z depletion could also have induced a DNA damage and stress response that could have triggered reactivation signals. Activation of p53 has been implicated in the initiation of the EBV lytic cycle (63, 64), and GO analysis of our RNAseq identified p53 pathway activation upon H2A.Z knockdown (Fig. 7B). We also observe a more significant perturbation of the cMyc and mTORC1 gene pathways, both of which have been implicated in the control of EBV reactivation (37, 65). H2A.Z has been found to regulate both MYC gene expression (66). This suggests that H2A.Z may be part of a more global program to control pathways important for EBV latency, including the control of BZLF1 and many cellular genes important for latency maintenance.

We also found that H2A.Z affected histone modifications on the EBV genome, including sites not bound directly by H2A.Z. Depletion of H2A.Z led to a loss of H3K27ac at Qp, as well as H4K20me3 at the inactive oriLyt region (Fig. 6). H2A.Z depletion did not lead to a loss of CTCF binding at multiple sites, suggesting that not all epigenetic features were stripped from the EBV genome. EBV lytic cycle is known to be regulated by multiple different control mechanisms, and the formation of repressive H3K20me3 at Zp and oriLyt are likely to maintain the latent state (67). Thus, it is possible H2A.Z may also regulate EBV latency through its effects on epigenetic patterning across the viral genome.

In conclusion, H2A.Z is an essential component of the EBV epigenome that is required for the stable maintenance of latency. H2A.Z colocalizes with EBNA1 and CTCF sites throughout the EBV genome and facilitates EBNA1 binding and function at *oriP* and Qp. H2A.Z depletion also disrupts many cellular gene transcripts, including those associated with myc and mTORC1 pathways. These studies further our understanding of the EBV epigenome and the role of histone variant H2A.Z in the control of viral latency.

## Material and Methods

### Cell lines

SNU719, LCL352, Mutu I, C666-1 cells were grown in RPMI1640 media supplemented with 10% fetal bovine serum, 1X Glutamax, and 100 μg/ml streptomycin, and 100U/ml penicillin, cells were incubated at 37°C with 5% CO2 in a humidified chamber. 293T (ATCC) cells were maintained in Dulbecco’s modified Eagle’s medium (DMEM) supplemented with 10% fetal bovine serum, 100 μg/ml streptomycin, and 100U/ml penicillin. Cells were cultured in an incubator set at 37 °C and 5% CO_2_.

### Lentiviral transduction

Lentiviruses were produced in HEK293T cells using envelope and packaging vectors pMD2.G and pSPAX2. The cells were co-transfected with pMD2.G and pSPAX2 and two vector-based shRNA constructs for H2AFZ (shH2A.Z-A and H2A.Z-B) and control shRNA (shControl), were generated in the pLKO.1 vector with the target sequence 5′-TTATCGCGCATATCACGCG-3′ (Table 1). After 48hrs post-transfection, supernatants were collected, then spun at 1000 x g for 15 mins to remove cellular debris, and then passed through a 0.45μM filter. Mutu I cells were resuspended in 10 ml lentivirus-containing supernatant and spun at 450 x g for 90 mins with 8μg/ml polybrene (Sigma-Aldrich, USA). The cells pellets were resuspended and incubated in fresh RPMI medium, after 48hrs, the cells were treated with 2 μg/ml puromycin. The RPMI medium with 2μg/ml puromycin was replaced every 2 to 3 days. The cells were collected after 7 days of puromycin selection and then subjected to further analyses.

**Table 1.**
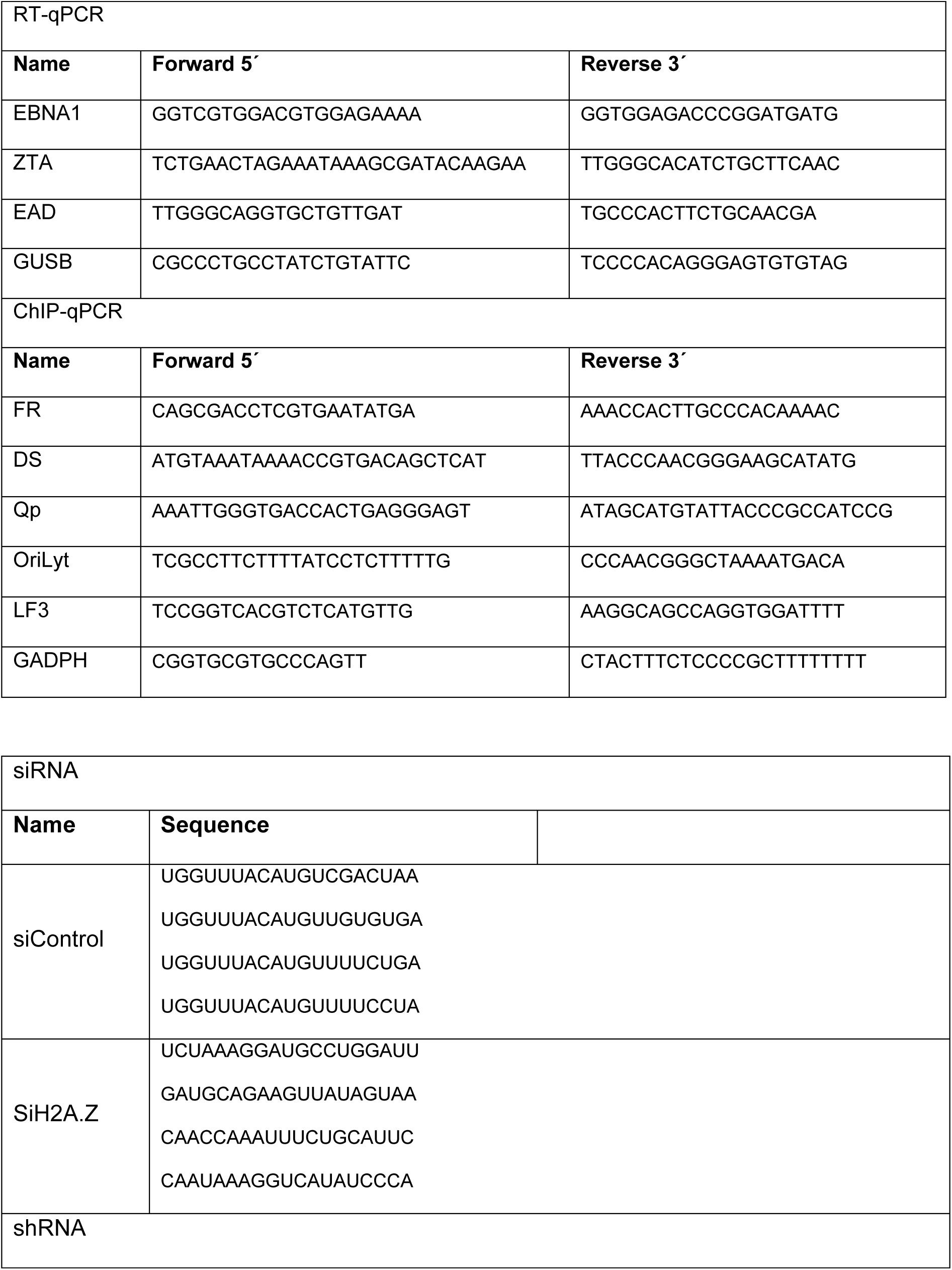
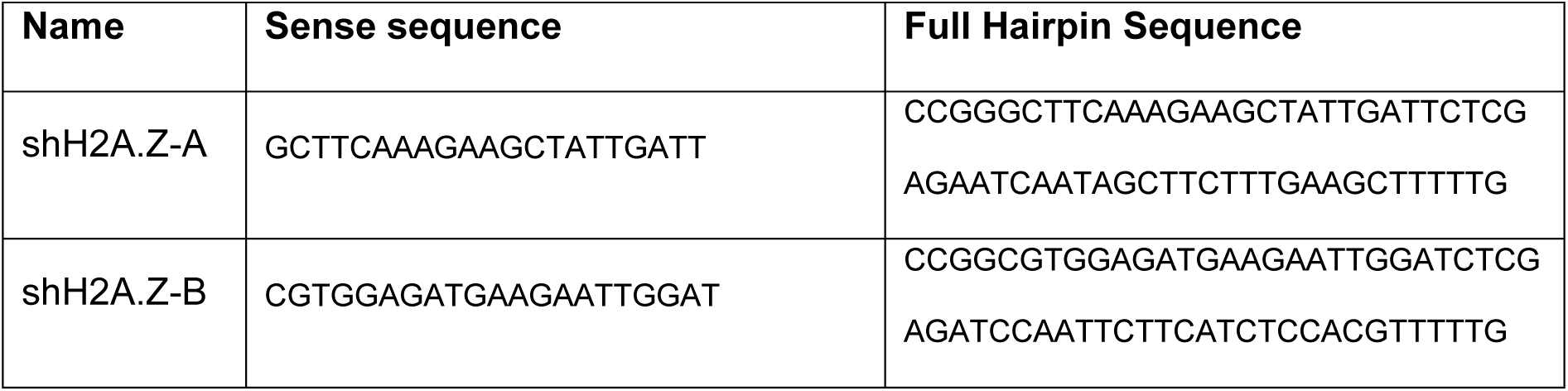
List of primers used.

### H2A.Z knockdown by siRNA

SNU179 cells were transfected with small interfering RNAs targeting siH2A.Z or siControl (Dharmacon, Lafayette, CO, USA) used DharmaFECT transfection reagents (Dharmacon, Lafayette, CO, USA). The cells were collected after 7 days of puromycin selection and then subjected to further analyses.

### RNA extraction and cDNA synthesis

Total RNA was isolated from cells using the RNeasy Mini kit (Qiagen, Hilden, Germany), according to the manufacturer’s protocol. RNA was resuspended in 30 μL of RNAse-free H_2_O, which was treated with the DNAse Free DNA removal kit (Thermo Fisher Scientific, Waltham, MA, USA) and quantified. Reverse transcription was carried out on equal amounts of DNase-treated RNA using SuperScriptIV reverse transcriptase kit (Invitrogen), following the manufacturer’s instructions. qPCR was performed with Power SYBR green 2X PCR master (ThermoFisher Scientific, Waltham, MA, USA) mix with specific primers. The expression of EBV genes were verified by RT-qPCR using specific primers (Table 1), and normalized to cellular GUSB gene expression.

### RNA-seq

Total RNA was isolated from each condition using the RNeasy Mini kit (Qiagen, Hilden, Germany), according to the manufacturer’s protocol. RNA was resuspended in 30 μL of RNAse-free H_2_O, which was treated with the DNAse DNA-Free DNA removal kit (Thermo Fisher Scientific, Waltham, MA, USA) and quantified by Qubit (Thermo Fisher Scientific, Waltham, MA, USA). RNA samples were either used for downstream RT-qPCR or submitted to the Wistar Institute genomics core facility for RNA quality control and sequencing library preparation using the SENSE mRNA-Seq Library Prep Kit V2 (Lexogen) to generate Illumina-compatible sequencing libraries according to the manufacturer’s instructions. Paired-end reads of 75 bp were obtained using a Illumina HiSeq 2500 sequencer. RNA-seq data was aligned using Bowtie2 (68) against hg19 version of the Human genome and all unaligned reads were then aligned against NC_007605.1 version of EBV genome and RSEM v1.2.12 software (69) was used to estimate raw read counts and RPKM for Human and EBV genes.

### Western blot

Cells were collected and protein extracts obtained by adding Laemmli sample buffer. Equal amounts of proteins were resolved in 8–16% Novex Tris-Glycine gels (Invitrogen), and then transferred onto a Nitrocellulose membrane (BIO-RAD-1620112). The membrane was blocked using 8% milk in TBS/0.01% Tween buffer and then incubated with the indicated antibodies. Primary antibodies were prepared in TBS/0.01% Tween buffer as follows: rabbit monoclonal anti-H2A.Z (Abcam-188314), rabbit polyclonal anti-EBNA1 (Pocono Rabbit Farm custom preparation), Anti-β-Actin (Sigma-A3854), rabbit polyclonal anti-ZTA (Pocono Rabbit Farm custom preparation), mouse monoclonal anti-EAD (Millipore-MAB8186). Membranes were washed three times with TBS/0.01% Tween buffer and incubated with HRP coupled secondary anti-mouse or anti-rabbit antibodies (Bio-Rad,). Finally, Proteins were visualized using the Immobilon Forte (Merck Millipore-WBLUF0500) according to the manufacturer’s instructions.

### Relative DNA copy number assay

Cells (1 × 10^6^ cells per sample) were collected and resuspended in 100 μl of SDS lysis buffer (1% SDS, 10 mM EDTA, 50 mM Tris [pH 8.0]). After 20 cycles of sonication, immunoprecipitation dilution buffer (0.01% SDS, 1.1% Triton X-100, 1.2 mM EDTA, 16.7 mM Tris [pH 8.0], 167 mM NaCl) was added to 1 ml, followed by incubation with proteinase K for 2 to 3 h at 50°C. Three hundred microliters of lysate were then removed and subjected to phenol-chloroform extraction and ethanol precipitation. Precipitated DNA was assayed by RT-qPCR. Relative EBV copy numbers were determined using primers (Table1) for different regions of EBV and normalized by the cellular DNA signal for the GAPDH or actin gene locus.

### Chromatin immunoprecipitation (ChIP)

For each ChIP assay, 1×10^6^ cells were crosslinked with 1% formaldehyde at room temperature for 10 min and the reaction was quenched with 0.125 M glycine for 5 min. Cells were lysed in 1 ml SDS lysis buffer (1% SDS, 10 mM EDTA, and 50 mM Tris-HCl, pH 8.0) containing protease inhibitor cocktails (Sigma-Aldrich), and kept on ice for 10 min. Lysates were sonicated with a Diagenode Bioruptor, cleared by centrifugation to remove insoluble materials, and diluted 10-fold into IP Buffer (0.01% SDS, 1.1% Triton X-100, 1.2mM EDTA, 16.7mM Tris pH 8.0, 167mM NaCl, 1 mM PMSF, and protease inhibitors cocktail). For each IP, 5 μg of anti-H2A.Z (Abcam-188314), Anti-EBNA1 (Pocono Rabbit Farm custom preparation), Anti-H3K27ac (Abcam, ab4729), Anti-H4K20me3 (Invitrogen,703863), Anti-H3K4me3 (Sigma-Aldrich, 07-473), Anti-H3K27me3 (Abcam,ab195477) or IgG (Cell Signaling, 2729) was added and rotated at 4°C overnight. Preblocked protein A sepharose (GE Healthcare, 17-0780-01/17-0618-01) was added to each IP reaction for additional 2 to 3 h incubation at 4°C. Beads were washed sequentially with low salt, high salt, LiCl, and TE buffer and then eluted with elution buffer (1% SDS, 0.1M NaHCO3). The eluates were then incubated at 65°C overnight to reverse cross-linking, followed by addition of Proteinase K at a final concentration of 100 μg/ml at 50°C for 2 hrs. ChIP DNA was purified by Quick PCR Purification Kit (Life Technologies) following the manufacturer’s instruction. ChIP DNA was assayed by qPCR using primers specific for indicated regions (Table 1). The relative enrichment was calculated as a percentage of input.

### H2A.Z and EBNA1 Chromatin immunoprecipitation sequencing (ChIP-seq)

Chromatin immunoprecipitation with next-generation sequencing (ChIP-seq) was performed as previously described. Briefly, 25×10^6^ cells per immunoprecipitation were collected and fixed with 1% formaldehyde for 15 min and then quenched with 0.25 M glycine for 5 min on ice. After 3 washes with 1× phosphate-buffered saline (PBS), pellets were resuspended in 10 mL each of a series of three lysis buffers before fragmentation in a Covaris ME220 sonicator (peak power 75, duty factor 25, cycles/burst 1,000, average power 18.8, time 720 s) to generate chromatin fragments roughly 200–500 bp in size as determined by DNA gel electrophoresis. Chromatin was centrifuged to clear debris and a 1:20 of this cleared chromatin was kept as standard input for comparison against immunoprecipitations. Chromatin was incubated by rotating at 4°C for 1 h with 25µg anti-H2A.Z (Abcam-188314) or affinity purified rabbit anti-EBNA1 (Pocono Rabbit Farms). Chromatin–antibody complexes were precipitated using a 50 µL of Dynabeads Protein A (ThermoFisher, product No. 10001D) incubated by rotating at 4°C overnight. Beads were then washed five times with RIPA buffer and TE buffer. Crosslinks were reversed by 65°C incubation with proteinase K followed by RNase A treatment. DNA was purified by Quick PCR Purification Kit (Life Technologies) following the manufacturer’s instruction. Libraries for sequencing were made using the NEBNext Ultra II DNA Library Prep Kit (New England Biolabs, product No. E7103) and sequenced on the NextSeq 500 (Illumina).

### ChIP-seq analysis

Reads were mapped against the human genome and human gammaherpesvirus 4 (HHV4) NC_007605.1 genome assembly using bowtie2 (70). We used MACS2 software packages to call reads enrichment in pull-down samples compared to input samples as peaks (71, 72). Analysis of peak distribution under differentiated conditions was performed with the bedtools software package (73) for genome arithmetic, and for data visualization we used deepTools (74). ChIP-seq data were deposited for public access at Gene Expression Omnibus (GEO accession number:xxx)

### RNAseq

RNAseq data was aligned using *STAR* (75) against hg19 version of the Human genome and all unaligned reads were then aligned against NC_007605.1 version of EBV genome and RSEM v1.2.12 software (24) was used to estimate raw read counts and RPKM for Human and EBV genes. Differential gene expression was obtained using DESeq2 R package (76). Statistically significant differences in gene expression were determined using a threshold of FDR < 0.05. Heatmaps were generated using matplotlib for Seaborn Python data visualization. Gene set enrichment analysis was performed using QIAGEN’s Ingenuity Pathway Analysis software (IPA; QIAGEN, Redwood City, CA; www.qiagen.com/ingenuity). RNAseq data were deposited for public access at Gene Expression Omnibus (GEO accession number:xxx)

### Statistical analysis

Data showing the effects of different assays are presented as mean ± SD. p was calculated by Anova and Student’s t-test analysis. Significant differences were accepted with p ≤ 0.05, as indicated.

## Acknowledgements

We thank members of the Wistar Cancer Center Cores in Genomics and Bioinformatics for their excellent technical support. This work was supported by grants from NIH DE017336, P01 CA269043-01A1, R01 CA140652, R01 CA093606 (PML), CA2059171-02S1 (LJCM) and P30 Cancer Center Support Grant P30 CA010815 to the Wistar Institute (D. Altieri).

